# Barriers Broken: Genetic Swamping in Restored Brook Trout Populations

**DOI:** 10.1101/2025.06.25.661651

**Authors:** Rebecca J. Smith, Matt A. Kulp, Benjamin M. Fitzpatrick

## Abstract

Scientists use reintroductions to restore native species to their historical ranges but sometimes can overlook the effects of dispersal on the genetic structure of restored populations. In the Great Smoky Mountains National Park (GRSM), an ambitious reintroduction strategy aims to restore native Brook Trout (*Salvelinus fontinalis)* populations, which have lost over 75% of their historical habitat within the park. Each restoration requires removing non-native trout and selecting restoration streams with natural barriers to prevent recolonization. Multiple Brook Trout populations are mixed in restoration sites to maximize genetic diversity and to avoid depletion of any single source population. However, a miscommunication during a reintroduction in Anthony Creek, which used three source populations for restoration, inadvertently created an in-situ population genetics study. One segment of the restoration site received fish from all three source populations. However, technicians also unintentionally placed fish from just one source upstream of a natural barrier. Unbeknownst to management, fish from this separate translocation dispersed downstream, potentially altering the desired genetic diversity of the restored population. This study characterizes the genetic changes caused by this unidirectional dispersal. Population genetics theory predicts that such movement leads to genetic swamping. Here, we use genetic and population density data to confirm the directionality of dispersal, estimate the rate of genetic swamping, and assess alternative mitigation strategies. Our results indicate that it is already too late for assisted migration within the restored stream to achieve the intended genetic diversity. Instead introducing additional fish above the second natural barrier would be necessary to equalize the contribution of all three source populations. Understanding the interplay between dispersal behavior and genetic structure is crucial for planning reintroductions and refining conservation strategies for this iconic species.

## Introduction

Genetic diversity enhances population resilience by increasing the likelihood that some individuals carry traits beneficial for survival in changing environments (1–3). Increasingly negative impacts from climate change, environmental disasters, and habitat fragmentation disrupt the genetic variation of wildlife and fish populations, putting species at risk by reducing genetic diversity and potentially hindering adaptation to environmental changes (4,5). Therefore, genetic management strategies emphasize the intentional maintenance or enhancement of genetic diversity (4).

Species reintroductions are an effective strategy to enhance genetic diversity and promote population resilience. Reintroductions work to restore populations to areas where they have been locally extirpated (6,7). Because threatened species generally exist in small, isolated populations, managers may use multiple sources for a single reintroduction site to prevent the depletion of any one source population (3,8). Mixing source populations is also intended to enhance genetic diversity, giving the restored population a greater potential to adapt to novel conditions and aiming for equal genetic contributions from each source while maximizing overall genetic diversity (9).

Brook Trout (*Salvelinus fontinalis*), a native species of the Southern Appalachian Mountains, has experienced significant population declines due to habitat loss and competition with nonnative salmonids (10)). In the Great Smoky Mountains National Park (GRSM), a Brook Trout reintroduction program restores the species to their historical range, where they have been locally extirpated (11,12). The GRSM Brook Trout reintroduction program utilizes multiple sources for each reintroduction, aiming to maximize genetic diversity and equalize genetic contributions from each source while avoiding the depletion of any single source. Prior to reintroduction, nonnative trout (Rainbow Trout, *Oncorhynchus mykiss*, and Brown Trout *Salmo trutta*) are removed from the targeted stream sections (11). A barrier, whether natural or artificial, is an essential component of these reintroduction efforts to prevent the recolonization of nonnative trout. However, such barriers also prevent upstream dispersal of Brook Trout and can inhibit the connectivity and adaptability of reintroduced populations.

Anthony Creek (Fig 1), one of the park’s restoration sites, contains multiple barriers restricting upstream fish movement, and translocated fish from multiple sources were released unequally among stream segments. In 2017, managers first stocked Anthony Creek with fish sourced from Bunches Creek, distributing them across two sections separated by a natural cascade barrier, which is likely to prevent upstream movement. Crews placed about half of the translocated fish above the cascade on the upper left fork while stocking the remaining fish in the section between the upper left fork barrier and a lower main stem barrier (an old mill dam) that prevents recolonization by non-native Rainbow Trout. The following year, Brook Trout from two additional source populations, Deep Creek and Sahlee Creek, were introduced into the intended mainstem restoration section between the upper left fork barrier and the lower barrier that prevents Rainbow Trout recolonization (Fig. 1-2). If fish can disperse downstream but not upstream, then unidirectional gene flow could result in genetic swamping (4,13). Specifically, if fish from Bunches Creek, located above the left fork barrier, continue to reproduce and disperse downstream, their genetic influence may overwhelm that of the Deep Creek and Sahlee Creek fish. This could reduce intended genetic diversity from all three source populations within the restored population between the two barriers, ultimately undermining the goals of the reintroduction effort.

**Fig 1:**
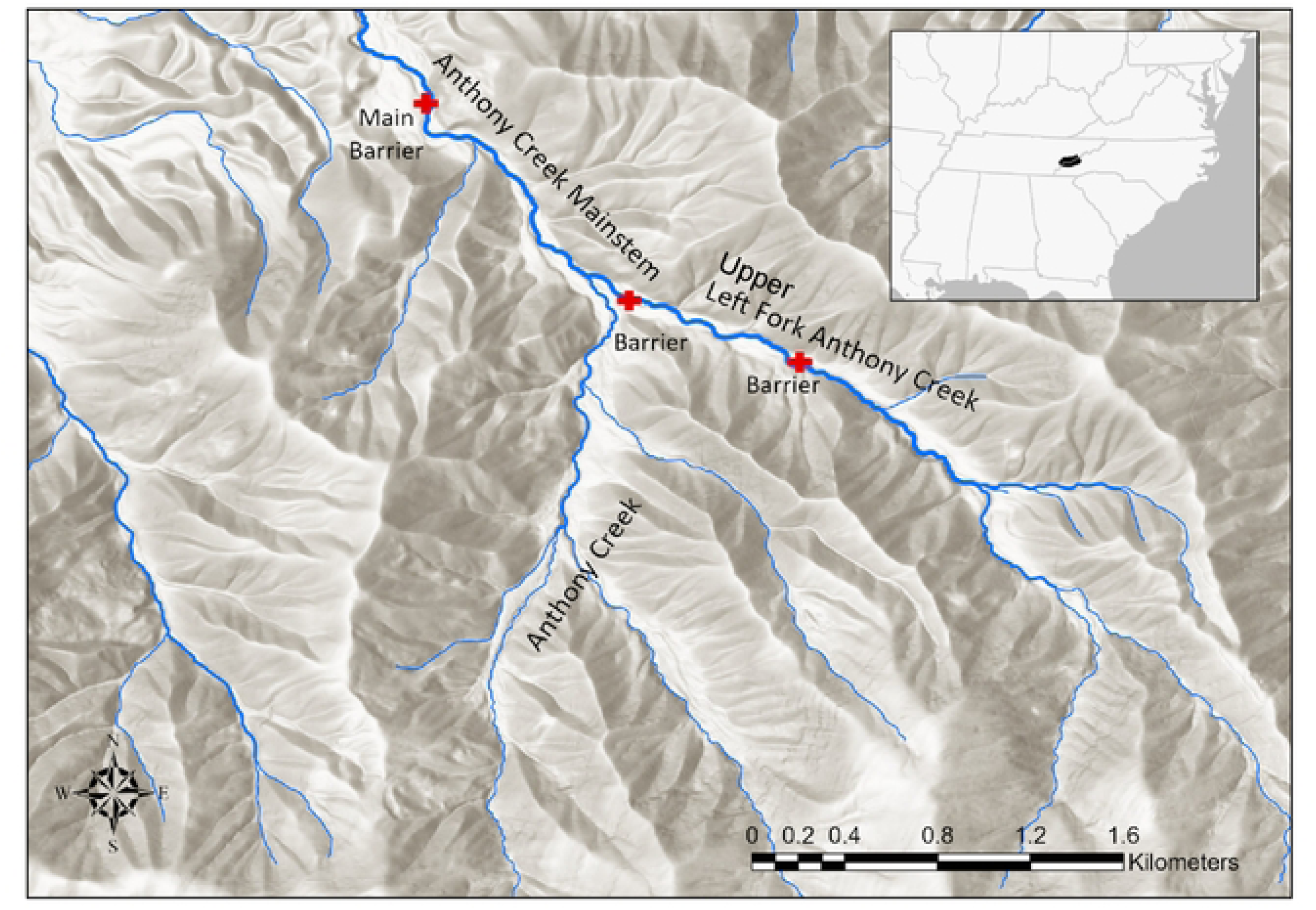
A map of Anthony Creek restoration site within The Great Smoky Mountains National Park. Red crosses mark the locations of in-stream barriers, and line width represents stream order. Mainstem and Left Fork Anthony Creek are both third order streams. GRSM is located in the Southeastern United States, spanning eastern Tennessee and western North Carolina.

**Fig 2:**
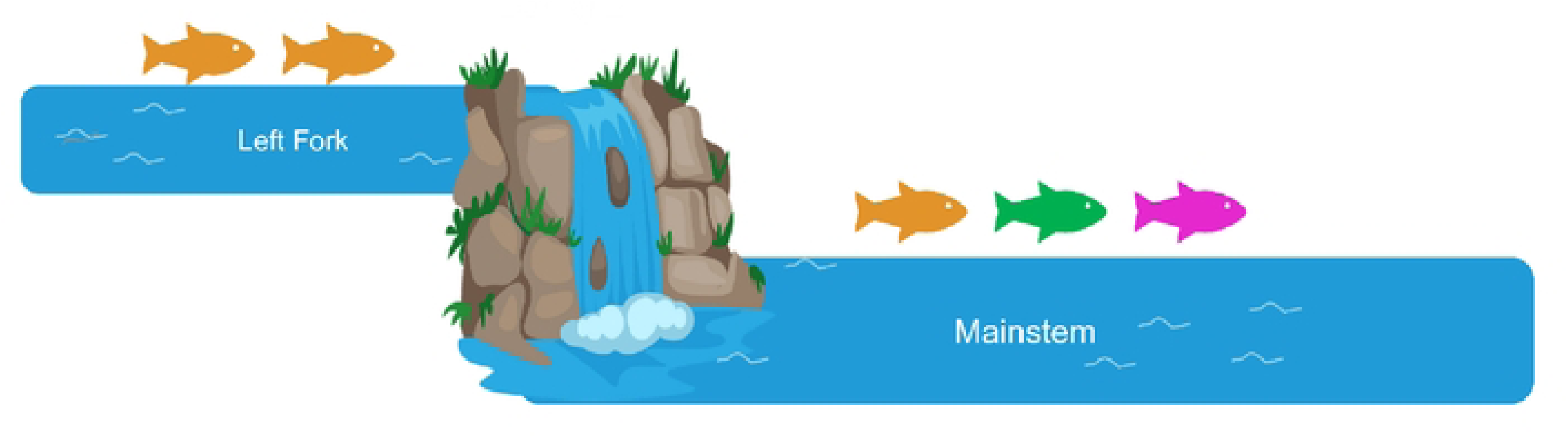
A schematic illustration of the translocation and stocking scenario of Anthony Creek. Fish from Bunches Creek (orange) were stocked in the Left Fork of Anthony Creek. Fish from Bunches (orange), Deep Creek (green), and Sahlee Creek (pink) were stocked in the mainstem, with a barrier preventing upstream dispersal between the two segments. Part of figure was created with Biorender.com

In this study we evaluate how unidirectional dispersal has affected the genetic diversity and structure of the restored population. We used ancestry analysis to confirm there is unidirectional dispersal and genetic swamping in the mainstem of Anthony Creek. We used stochastic simulation modeling to estimate the rate of unidirectional immigration. Next, we forecasted the ancestry proportions to the present day and used predictive models to estimate how many fish from Deep and Sahlee Creeks need to be added above the barrier to equalize the intended genetic contributions from each source.

## Methods

### Study System

Anthony Creek (Figs 1-2) did not have extant populations of Brook Trout before reintroductions, and nonnative trout were removed before translocation (14). Based on prior GRSM Brook Trout translocation experiences, the source populations are selected from within the same major watershed (Little Tennessee River) as the restoration area. The management goal is to balance the preservation of source population numbers while achieving a target translocation density of 125-150 Brook Trout per kilometer at the restoration site (11,12,14).

Anthony Creek was initially stocked with 269 individuals from Bunches Creek, with the translocated fish distributed between the left fork (n = 135) and the mainstem (n =134) of Anthony Creek. The fish were intended to be placed solely within the mainstem but were instead distributed between both areas due to miscommunication between field crews. In 2018, fish were translocated from Deep Creek (n = 135) and Sahlee Creek (n = 102) into the mainstem of Anthony Creek (Table 1). Management was unaware that fish had been stocked in the upper left fork until 2022. During that summer, we collected DNA samples from the three source populations, as well as from both the left fork and mainstem of Anthony Creek.

**Table 1:** Overview of populations included in the study, with the corresponding number of samples (N) collected from each population and the year of sampling.

|  |  |  |
| --- | --- | --- |
| Population | Sampled Year | N |
| Anthony Creek (left fork) | 2022 | 29 |
| Anthony Creek (mainstem) | 2022 | 64 |
| Bunches Creek | 2022 | 100 |
| Deep Creek | 2022 | 98 |
| Sahlee Creek | 2022 | 60 |

Since the time of sampling, the composition of the Anthony Creek restoration has been made more complicated by an additional translocation of Brook Trout to a third stream segment. This segment is above the left fork segment and separated from it by another natural cascade. The segment was stocked with even mixture of fish from Bunches Creek and Louie Camp Creek.

Thus, consideration of future actions will have to account for this most upstream population. Here we use our data from the left fork and main stem to test the hypothesis of unidirectional dispersal and estimate the rate of downstream immigration to understand the consequences of the spatially structured translocation history, and the feasibility of hypothetical interventions to maximize long-term genetic diversity.

### Sample Data & Processing

We captured live fish during regularly scheduled abundance surveys and removed adipose fin clips using sterile scissors prior to re-releasing fish alive within 100 meters of capture (11,15). The fin clips are collected by National Park Service staff for management purposes and are excluded from coverage under the Institutional Animal Care and Use Committee (IACUC) or Animal Welfare Act. All procedures complied with the American Fisheries Society Guidelines for the use of fish in research (16) and were approved under Scientific Research and Collecting Permit GRSM-SCI-2158. We extracted DNA using the Qiagen DNAeasy blood & tissue kits.

The Idaho Department of Fish and Game Eagle Genetics Lab performed the sequencing using a targeted sequencing panel of 304 loci developed by the lab in collaboration with the Columbia River Inter-Tribal Fish Commission for Brook Trout. The panel includes loci from (17) and additional loci identified through restriction site-associated DNA sequencing (Columbia River Inter-Tribal Fish Commission, unpublished data; IDFG, unpublished data). Sample preparation and sequencing followed Genotyping in Thousands Sequencing (GTseq) protocols (18). The Idaho Department of Fish and Game originally designed the targeted sequencing panel to score predefined SNPs in the Columbia River. However, we used the complete sequence data to discover and score all SNPs in our GRSM samples. IDFG filtered out low-quality reads and adapter contamination. We mapped the filtered and demultiplexed 79 base pair reads to the reference sequences using BWA mem (19). We called variants using bcftools mpileup in the samtools suite (20), excluding those with a quality score below 200 or a depth below 20. Next, we examined the resulting VCF file manually in R (21) and tailored a quality control protocol to include additional filtering steps specific to our dataset as follows.

Initial filtering steps included the removal of three individuals identified as Rainbow Trout (*Oncorhynchus mykiss)*. Indels and invariant SNPs were excluded from the dataset using the extract.indels and is.polymorphic functions from the vcfR package (22), retaining only polymorphic SNP loci. We removed individuals with >5% missing genotypes. After these filtering steps, fewer than 20% of SNPs had still any missing data in any individual. To ensure a final dataset with no missing data we excluded all of them to produce a data set with no missing data across all individuals. Collectively, the foregoing steps eliminated approximately half of the original 304 loci. The filtered VCF was converted to a genind object for multivariate analysis.

We visually inspected the data with PCA plots which revealed clustering patterns associated with sequencing plates, suggesting potential batch effects. To mitigate this, we implemented a DAPC- based feature selection approach using the snpzip function in the glmnet package (23) to identify loci strongly associated with plate identity. A total of 48 SNPs were flagged and removed, effectively eliminating batch structure in the data. We then selected the first SNP from each locus to preserve the assumption of independence for ancestry estimation. The final filtered dataset included high-confidence genotype data for 232 alleles across 116 SNP loci. All sequencing data are available online under NCBI BioProject ID PRJNA127654. Additional materials are provided in Supplemental File S1.

### Ancestry and Migration Rate Estimation

To determine the relative contribution of each source to the restored populations in Anthony Creek, we estimated ancestry proportions for each sample using HIest (24). We estimated source population allele frequencies from the source population samples and used these to estimate the ancestry proportions in Anthony Creek left fork and mainstem. The threeway function estimates the most likely set of three ancestry proportions for each individual given its genotype and the given source allele frequencies. We sampled only 5 years after mixing (the fourth breeding season). After only four breeding seasons, we expect a mixture of predominantly parental, F1, F2, and backcross genotypes because few if any F2 and backcrosses would have reached sexual maturity by the time of sampling (25). If the first F1s were spawned as soon as possible in 2018, some could have been sexually mature by 2020, and their offspring (F2 and backcrosses) could be sexually mature by the 2022 breeding season, after sample collection. It is extremely unlikely that we would have sampled F3s or later generation backcrosses. If there was no dispersal between restored stream segments, we would expect all samples from the left fork (above the cascade) to have 100% Bunches Creek ancestry, and samples from the mainstem to be a mixture with approximately 42% Bunches Creek ancestry, resembling the input proportions from each source. If there has been downstream dispersal, the proportion of Bunches Creek ancestry in the mainstem would be a function of the immigration rate and time (26):

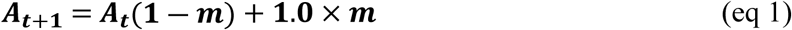

where *A_t_* is the Bunches ancestry proportion in the mainstem at time *t*, *m* is the immigration rate per time interval, and 1.0 is the ancestry proportion of immigrants. This equation models expected change in ancestry proportions from year to year, assuming a constant immigration fraction *m*. The general solution

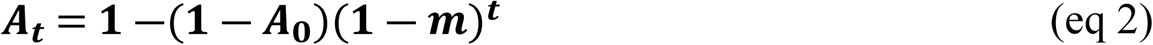

predicts the ancestry proportion in any year *t* as a function of the immigration rate and an initial ancestry proportion *A*_0_.

To estimate the range of downstream dispersal rates (*m*) consistent with the observed data, we conducted stochastic simulations of population growth and genetic drift according to the age-structured Brook Trout life history model (27). To reflect the known introduction history, we first simulated one year of reproduction starting with 135 Bunches Creek fish. In the second year (corresponding to 2018) we added 135 Deep Creek and 105 Sahlee genotypes. For each subsequent year of the simulation (up to 2022, the sampling year), we added pure Bunches Creek immigrants (implicitly from the left fork population) of varying age classes using a random draw from a multinomial distribution with size *N_t_m* (where *N_t_* is the total population size in the mainstem in year *t*) and probabilities determined from the stable age distribution determined from preliminary model runs (0.66 age 0, 0.21 for age 1, 0.08 for age 2, and 0.05 for age 3). We performed 10,000 replicates each for immigration rates between 0.1 and 0.25 in increments of 0.01. We then compared the distribution of simulated ancestry proportions for each immigration rate to the estimated value from our 2022 data. We defined an immigration rate as credible if the interquartile range of the simulated outcomes included the observed value of Bunches Creek ancestry in the main stem. R scripts for recreating these simulations are given in Supplemental file 2.

Because potential remediation could be undertaken in 2025 or later, we used the credible range of immigration rates to forecast the expected proportion of Bunches Creek ancestry in the mainstem population using equation 2. Thus, we use our estimated range of credible immigration rates to forecast expected future ancestry proportions in the mainstem with *A_0_* set equal to our estimated value from the 2022 samples.

### Remediation Strategies

Finally, we used these results to explore the feasibility of two strategies to recover desired genetic diversity. First, individuals from the genetically mixed mainstem population could be transported above the barrier cascade to modify the ancestry proportions of the left fork population. With continued subsequent one-way dispersal, the equilibrium ancestry proportion of the mainstem would converge on the new left fork ancestry proportion. For example, if *N_I_*immigrants with average ancestry *A_I_* could be transported and added to the population of *N_R_* residents in the left fork, the new ancestry proportion in the left fork would be eq 3:

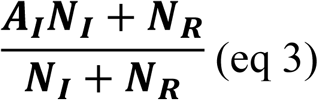

Alternatively, if the desired ancestry proportion is lower than what is possible given eq 3, then new fish could be translocated from the other original source populations (with zero Bunches Creek ancestry). In this scenario, because the ancestry proportion of the immigrants is *A_I_* = 0, the new (and expected equilibrium) ancestry proportion would be eq 4:

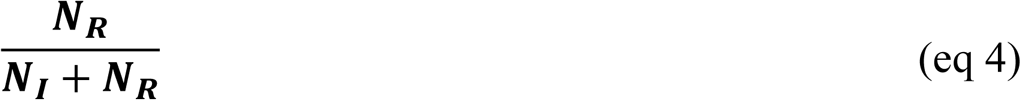

In either case, to determine the feasibility of translocations, we require an estimate of the total population size in the left fork (*N_R_*).

### Population Size

Along with the GRSM fisheries science team, we used population density estimates from annual three-pass depletion surveys to extrapolate a total population size for the left fork section. In 2019, 2020, 2023, and 2024 we conducted three-pass surveys of 100 m stream sections according to the electrofishing protocol described by Habera et al. (28), Kulp & Moore (14), and Kanno et al. (11). We used 6-10 mm bar mesh nets to block the lower and upper end of each 100 m site and captured every fish encountered on three sequential passes with backpack electrofishing units. We counted and identified each fish by species, recorded mass (g) and total length (mm), and retained them in holding cages outside of the site until the depletion survey was complete. We then released all fish back into the site. We estimated abundance of each size class using the Burnham maximum likelihood estimator in the Microfish 3.0 software (Deveneter et al., 1989).

During each survey, we measured stream width at 10 m intervals to estimate the surface area of each 100 m section at the time of the survey. We used these estimates to convert abundance estimates to densities (number of fish per m^2^). We then used the total stream distance of the left fork population (711m, bounded by cascades on both ends) and the average stream width to extrapolate a total stream area (average across years 2927 m^2^). We estimated total population size by multiplying this area by the average density. While more refined population size estimates may be possible (29), this simple extrapolation is sufficient for present purposes of demonstrating an approach for planning future translocations.

## Results

### Ancestry analysis and Dispersal patterns

Ancestry estimates in the left fork stream section, where only fish from Bunches Creek were released, were 100% Bunches Creek except for four individuals with a minority of their ancestry estimated as coming from either Sahlee or Deep Creek (Fig. 3). To evaluate whether those individuals are more likely to be backcrosses or mis-characterized because of estimation error, we used the classification functions in HIest (HIclass and HItest) to calculate the likelihoods of them being backcrosses, pure Bunches ancestry, or later generation backcrosses (24). Only one individual was more likely to be a backcross than a pure Bunches Creek fish, and the difference in likelihood was small (0.09 log-likelihood units). In other words, the SNP data were not sufficiently diagnostic to reliably distinguish pure Bunches ancestry from some possible backcrosses. These results are consistent with zero upstream dispersal, but do not rule out a very low rate.

**Fig 3:**
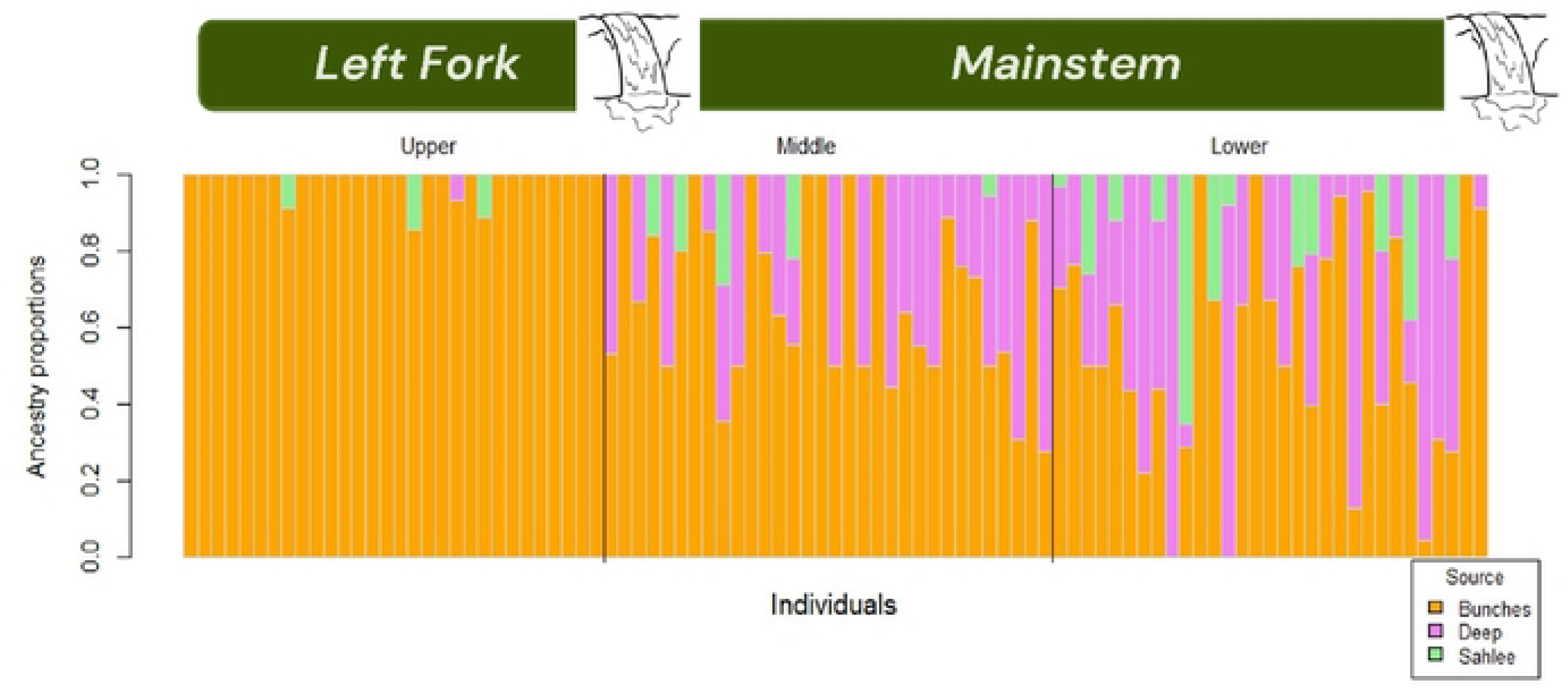
Ancestry proportions of individual Brook Trout in the restored population, estimated using HIest (Fitzpatrick 2012) and the estimated allele frequencies from the three source populations. Individual fish are represented by vertical bars arranged along the horizontal axis according which stream section they were captured in. The y-axis shows the proportion of genetic ancestry from each source population. Colors indicate the contribution of each source.

In the mainstem of Anthony Creek, Bunches Creek accounted for 63% of the total ancestry, while Deep Creek contributed 25%, and Sahlee Creek comprised 12% of the ancestry. Within the mainstem, average Bunches Creek ancestry was somewhat higher in fish collected closer to the upstream barrier (Fig. 3). While individual ancestry estimates are subject to error, we assume the population averages accurately reflect the biological processes of immigration and genetic drift.

### Migration Rate Estimation

To estimate a credible range of immigration rates from the left fork, above the barrier, to the mainstem below, we ran stochastic simulations until we found a range that fit the observed ancestry proportion of Bunches creek ancestry (63%) in the mainstem. For illustration, we declared an immigration rate credible if the interquartile range of the simulated outcomes included the observed value. Under this criterion, a suitable range of immigration rates is 13- 19% (Fig. 4), that is, 13-19% of the fish in the mainstem at any given time have come from the left fork in the previous year.

**Fig 4:**
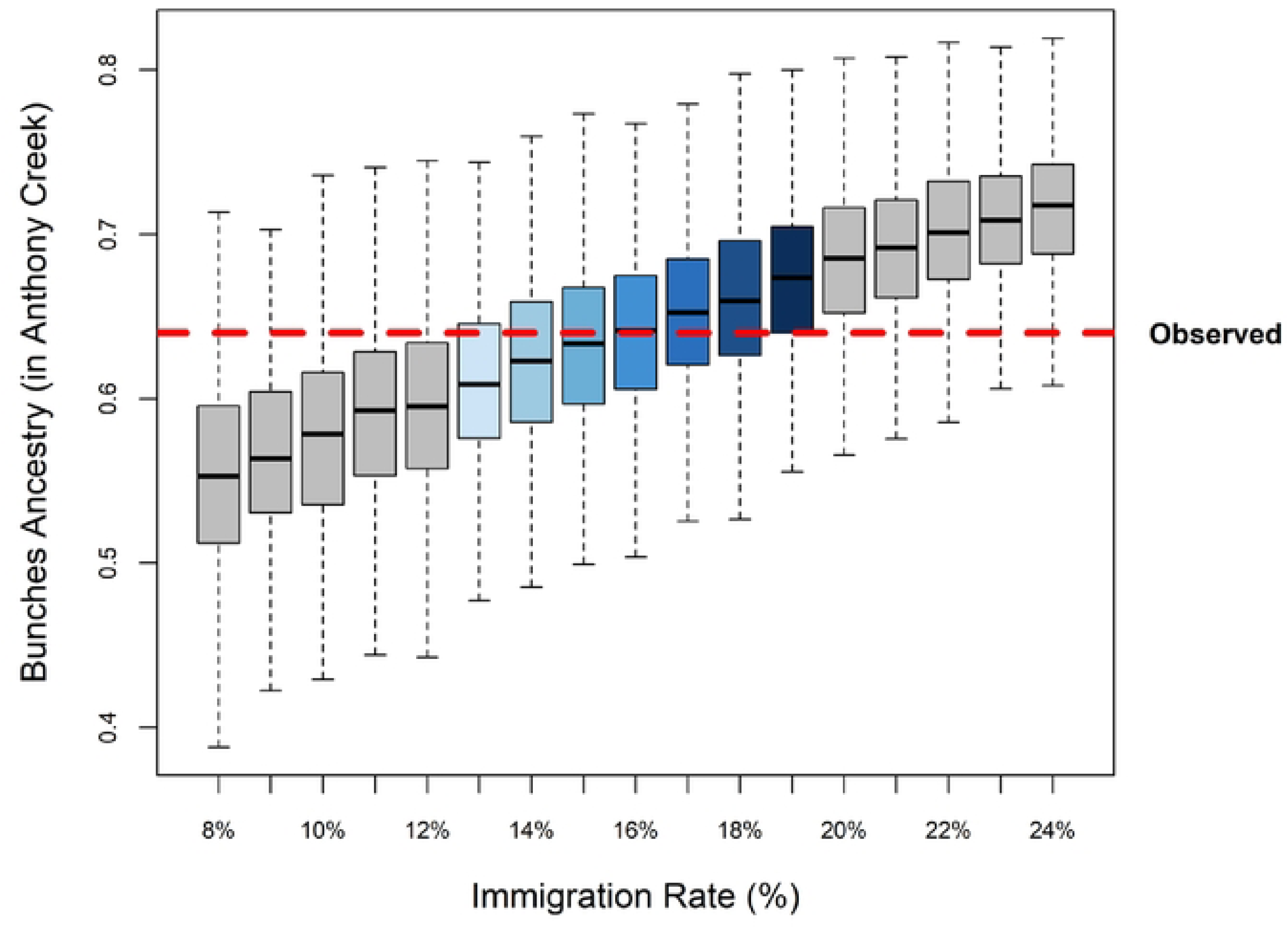
Box plot with range of immigration rates (%) on x axis and the ancestry proportion owing to Bunches Creek measured on the Y axis. The observed proportion in our sample is noted by the red dashed line. Boxes indicate the interquartile range and horizontal bars the median ancestry from xxx simulations of each immigration rate. Whiskers extend to the furthest point within 1.5 times the interquartile range from the box.

### Forecasting Bunches Creek Ancestry in Anthony Creek

We expect the proportion of Bunches’ ancestry in the mainstem to continue increasing without any intervention. In our five-year forecast, the mainstem of Anthony Creek is expected to reach 74-80% Bunches ancestry by 2025 and 78-87% by 2027, depending on the immigration rate (Fig 5).

**Fig 5:**
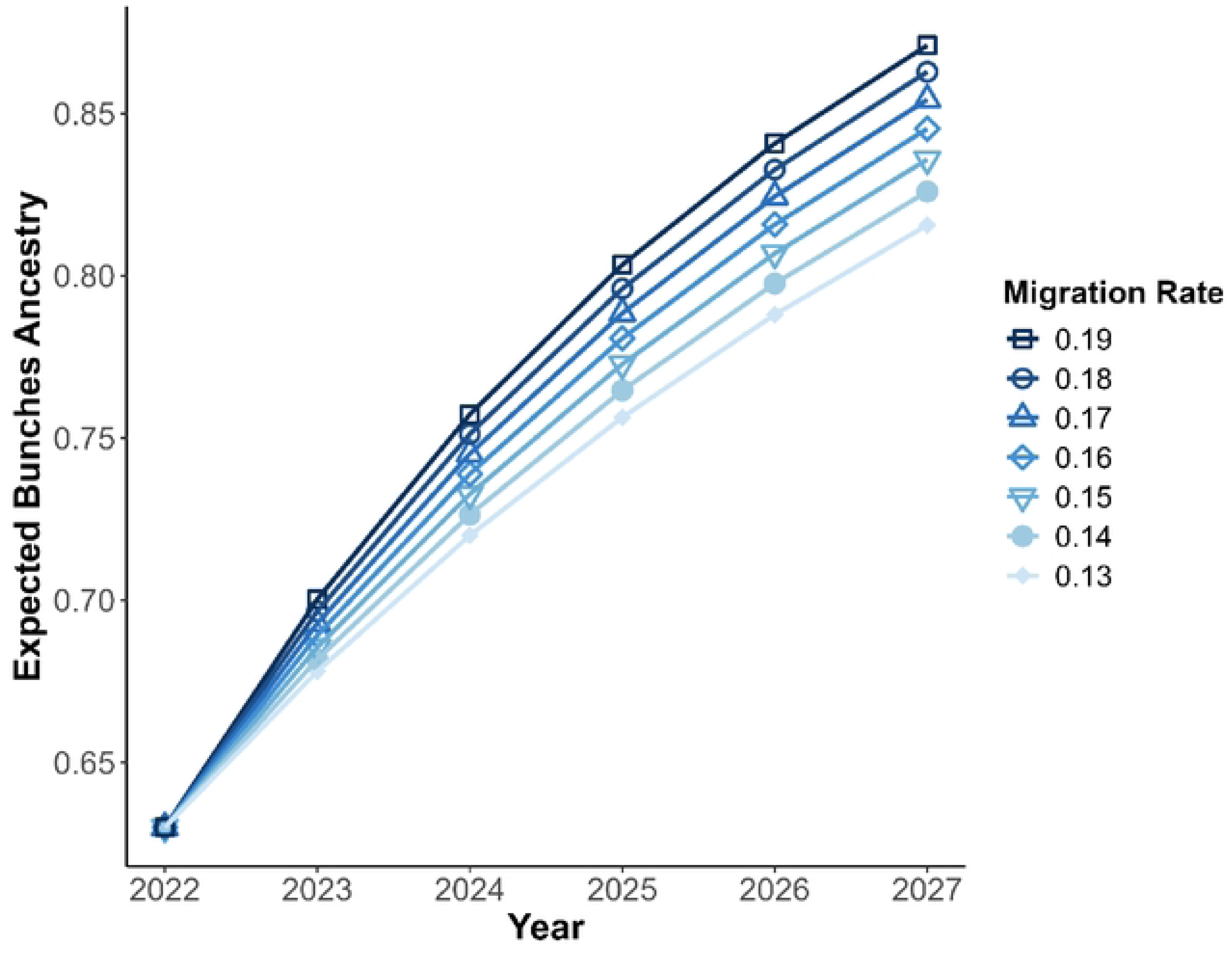
Forecasting Bunches Creek ancestry in the mainstem from the time of sample collection in 2022 to five years later in 2027. Using a range of migration rate estimates, we project that Bunches Creek ancestry will comprise 78-87% of the total ancestry by 2027.

### Population Size

Density estimates from three-pass depletion surveys in the Left Fork increased from 1.6 fish per 100m^2^ in 2019 and 2020 to 5.1 per 100m^2^ in 2023 and 12.5 per 100m^2^ in 2024 (see Supplemental File 3). The latter is comparable to the average density of 11.6 per 100m^2^ in restored Brook Trout populations reported by Kanno et al. (11). Extrapolating to the entire area of the Left fork (2927 m^2^), the total population size estimate for 2024 is 365 Brook Trout, and the average across years is 153.

### Remediation Strategies

Given the projected ancestry proportions in the mainstem, translocating fish from the mainstem to the left fork could stabilize the system at a high fraction of Bunches ancestry. For example, if we take the mainstem ancestry in 2025 as 80% (Fig. 6) and translocate enough fish to result in a 1:1 dilution, the resulting ancestry proportion would be 90%. If feasible, this would result in a doubling of the population density above the barrier, but relatively minor genetic change.

To achieve roughly 1/3 contribution from each source would require translocation of new fish from Deep Creek and Sahlee Creek. The most aggressive strategy of adding enough fish from each of Deep Creek and Sahlee Creek to instantly achieve roughly 1/3 contribution from each source, would triple the population density in the Left Fork, and would probably require removing too many fish from the source populations. We used the one-way immigration model (equations 1 and 2) to illustrate a more measured approach of translocating additional fish from Deep and Sahlee Creeks multiple years in a row (Table 2), to decrease the percentage of Bunches ancestry in the system. Table 2 lists the expected effects of adding specific numbers to an assumed constant population of 365 fish. Calculations are shown in Supplemental File 3.

**Table 2.** Expected results of sequential translocation of a fixed fraction of individuals into a population to achieve a desired ancestry proportion of 1/3 (Immigration Fraction, *m* = number of immigrants ÷ (number immigrants + number of residents)). The expected resident ancestry is expected to decay geometrically, *A_t_ =* (1 – *m*)*^t^*. The listed number per year is the total number of immigrants to be translocated into a resident population of 365 individuals. For example, if two sources were used, 45 from each source would sum to 90 immigrants and *m* = 90 ÷ (90 + 365) = 0.1978, resulting in 33.33% ancestry from each at the end of five years.

| Table 2. Expected results of sequential translocation of a fixed fraction of individuals into a population to achieve a desired ancestry proportion of 1/3 (Immigration Fraction, $m = \text{number of immigrants} \div (\text{number immigrants} + \text{number of residents})$). The expected resident ancestry is expected to decay geometrically, $A_t = (1 - m)^t$. The listed number per year is the total number of immigrants to be translocated into a resident population of 365 individuals. For example, if two sources were used, 45 from each source would sum to 90 immigrants and $m = 90 \div (90 + 365) = 0.1978$, resulting in 33.33% ancestry from each at the end of five years. | | | | | | |
| --- | --- | --- | --- | --- | --- | --- |
| Number Per Year | Immigration Fraction | Expected Resident Ancestry |  |  |  |  |
|  |  | Year 1 | Year 2 | Year 3 | Year 4 | Year 5 |
| 267.2 | 0.423 | 0.577 | 0.333 |  |  |  |
| 161.4 | 0.307 | 0.693 | 0.481 | 0.333 |  |  |
| 115.4 | 0.240 | 0.760 | 0.577 | 0.439 | 0.333 |  |
| 89.7 | 0.197 | 0.803 | 0.644 | 0.517 | 0.415 | 0.3333 |

## Discussion

Our results indicate that unidirectional dispersal has disrupted the intended equal genetic contributions from the three stocked source populations in the Anthony Creek restoration site. This one-way gene flow has resulted in the mainstem’s genetic ancestry and diversity being dominated by the Bunches Creek lineage. If one-way migration continues, the restored population may trend toward genetic homogenization in terms of ancestry. Equal genetic contribution is a common goal in mixed-source reintroductions, and moving fish solely from the mainstem into the left fork is unlikely to achieve this balance. To promote a more balanced representation of the three source populations stocked in Anthony Creek, additional translocations from Deep Creek and Sahlee Creek would be required. We have not tested whether any single source harbors selectively advantageous alleles or greater adaptive potential; without that information, an equal representation of all three sources is the best recommendation to ensure a diverse gene pool, increase the restored population’s capacity to adapt to environmental changes, and enhance long-term resilience.

Ancestry analysis confirms that upstream dispersal is negligible. However, distinguishing between strictly unidirectional migration and minimal movement in the opposite direction remains challenging. Our data could be consistent with rare upstream dispersal; however, these events do not significantly impact ancestry patterns or lead to equal contributions from each source. Therefore, even if limited upstream dispersal occurs, it would not meaningfully affect the conclusions or implications of this study.

Mainstem ancestry has shifted due to high immigration rates, increasing from the original 42% Bunches ancestry (2018) to a projected 74-80% (2025). Our estimated dispersal rate of 13-19% indicates that fish are moving at a rate that significantly influences downstream populations. This change highlights the importance of considering barriers and the directionality of gene flow in restoration efforts, especially when aiming for equal genetic contributions. These considerations are critical for enhancing genetic diversity in restored populations and planning for long-term resilience.

Achieving equal genetic contribution through translocating fish from the mainstem above the barrier into the left fork is unlikely to be the best solution, albeit least taxing on management and fish. It is not a good solution because Bunches Creek ancestry already dominates the mainstem’s genetic makeup. The most effective way to attain equal genetic contribution is to translocate additional fish from Deep and Sahlee Creeks into the left fork. Previous Brook Trout studies in GRSM (11) indicated steady state density average of approximately 12 fish per 100 m^2^, which is comparable to our estimates in 2024 (Supplemental file 1). This means the estimated 365 total fish in the left fork of Anthony Creek may be close to the local carrying capacity. Any additional translocations will increase population size and may push the site over what Anthony Creek can support. Therefore, it may be best to stagger translocation events over time to minimize demographic stress.

However, the situation is more complex. After our sampling, additional fish from a fourth source, Bearwallow Branch, were introduced above the upper left fork barrier to help increase fish abundance in Anthony Creek. As a result, the upper left fork population is now approximately 50% Bunches and 50% Bearwallow Branch. This may increase the overall immigration rate into the mainstem, but any effort to manage genetic diversity in this system should prioritize the uppermost population. If maintaining Deep and Sahlee genetic contributions across the system is a goal, additional translocations to the uppermost reaches would be necessary. This approach would require careful consideration of the current population size, carrying capacity, and the number of fish that could be added without exceeding these ecological constraints.

Strategic translocation planning is critical for maximizing genetic diversity and population stability. Considerations such as carrying capacity and whether translocations should occur in a single event or be staggered over multiple years can influence restoration success. Previous research on multi-stocked restorations in GRSM has shown that, in the absence of barriers, complete admixture can occur between sources (27). However, one-way migration can disrupt this balance by eroding diversity and favoring alleles from upstream populations.

The genetic consequences of unidirectional dispersal are particularly concerning in the context of local adaptation. Natural selection can counteract genetic swamping if the selection coefficient is greater than the immigration rate (31,32). However, given the high immigration rates inferred in our study, only exceptionally strong selection would be capable of retaining potentially advantageous alleles from Deep Creek and Sahlee Creek. Moderately advantageous alleles, if present in the mainstem population, might be preserved by translocating fish from the mainstem to the left fork (above the barrier), where selection would be expected to increase their frequencies even if the overall contribution of Deep Creek and Sahlee Creek ancestry remains low. These dynamics underscore the importance of considering both gene flow and selection pressures when designing conservation strategies.

These findings have broad implications for conservation efforts aimed at addressing habitat fragmentation, managing invasive species, and guiding reintroduction programs. Asymmetric dispersal behavior can significantly shape genetic composition in populations, potentially steering them toward more desirable outcomes. For example, stocking hatchery-reared fish has historically been a common practice to replace locally extinct populations or supplement existing ones for angling purposes (33). While modern conservation practices favor using native Brook Trout populations for stocking, remnant hatchery Brook Trout populations persist on the landscape (34). One-way migration could potentially be strategically used to remove non-native, hatchery-reared fish, as continued gene flow from native populations could eventually swamp out unwanted hatchery genetics.

Reintroduction efforts remain critical for the conservation of many species. Applying genetic and evolutionary theory in practice can enhance the likelihood of creating resilient, self-sustaining populations. Thus, understanding gene flow dynamics is essential for refining restoration strategies and minimizing unintended genetic consequences in freshwater fish and other species facing similar ecological challenges.

## Author Contributions

R. J. Smith and B. M. Fitzpatrick conceived and designed the study. M. Kulp coordinated sample collection dates, and all authors participated in field sampling. R. J. Smith conducted the laboratory work to prepare samples for sequencing. R.J. Smith and B. M. Fitzpatrick contributed to data analysis and wrote the manuscript, and M. Kulp contributed provided edits and revisions.

## Acknowledgements

We thank Idaho Fish & Game’s Eagle Genetics Lab for their support with preliminary sequencing runs to test the Brook Trout panel with our samples, as well as for their contributions to subsequent sequencing efforts. Special thanks to Caleb Abrahmson and the rest of the Great Smoky Mountains National Park Fish Science Team for their invaluable assistance with sample collection.

